# Development and optimization of a germination assay and long-term storage for *Cannabis sativa* pollen

**DOI:** 10.1101/2020.03.19.999367

**Authors:** Daniel Gaudet, Narendra Singh Yadav, Aleksei Sorokin, Andrii Bilichak, Igor Kovalchuk

## Abstract

Pollen viability and storage is of great interest to cannabis breeders and researchers to maintain desirable germplasm for future use in breeding or for biotechnological and gene editing applications. Here, we report a simple and efficient cryopreservation method for long-term storage of *Cannabis sativa* pollen. Additionally, we have deciphered the bicellular nature of cannabis pollen using DAPI staining. We have also standardized a pollen germination assay to assess the viability of cannabis pollen, and found pollen collected from different principal growth stages exhibits different longevity. Finally, we developed a long-term storage method which includes pollen combination with baked whole wheat flower and desiccation under vacuum for cryopreservation. By using this method, we were able to maintain germination viability in liquid nitrogen after 4 months, suggesting potentially indefinite preservation of cannabis pollen.

## Introduction

Cannabis or hemp (*Cannabis sativa* L.) is an annual, primarily dioecious flowering plant. The center of origin is in Central Asia, and it has been bred for thousand of years for a variety of traits including fiber, oil, seed and drug use (Piluzza et al., 2013). Cannabis is a diploid plant (2n=20), males are defined by heterogametic chromosomes (XY) with homogametic (XX) conferring the female phenotype. Male plants produce flowers containing stamens producing pollen whereas female plants develop ovaries that produce seed following pollination. Female inflorescences are characterized by secretory hairs known as glandular trichomes which produce a resinous mix of cannabinoids and aromatic compounds that are valued for both medical therapeutics and recreational effects (Chandra and Lata, 2017).

Pollen viability is of great interest to breeders and researchers alike. Breeding projects may wish to store pollen for extended periods of time, where high value genetic material may be stored for future use or for biotechnological and gene editing applications, which requires a quick and effective method of determining pollen viability (Zottini et al., 1997; Engelmann and Takagi, 2000; Choudhary et al., 2014). Fluorescent stains such as fluorescein diacetate (FDA) or fluorochromatic reaction test (FCR) have been previously reported for assessing pollen viability in cannabis (Zottini et al., 1997; Choudhary et al., 2014). Viability is not always correlated with germination, as pollen may retain the ability to metabolize, while losing it’s ability to germinate (Engelmann and Takagi, 2000). To better assess germination, we established a pollen germination assay (PGA) to estimate germination rates. We also adapted a DAPI stain to visualize pollen pre- and post-germination, and to establish whether *Cannabis sativa* was a bicellular or tricellular species, which to our knowledge has not been reported in the literature. Approximately 30% of angiosperms are known to be tricellular, with the male gametophyte sexually immature at the time of anthesis (Williams et al., 2014). We also used the PGA to test how storage and timing of pollen collection could influence germination rates. Pollen germination rates were assessed over a period when stored at 4°C from males at different stages of floral development. Finally, we developed a simple procedure for the long-term storage of cannabis pollen using desiccation with baked whole wheat flower followed by cryopreservation which potentially maintain long-term viability of pollen for future use.

## Materials and Methods

### Plant material and growth conditions

*Cannabis sativa* plants (strain name “Spice”, THC dominant) were grown under full spectrum 300- Watt LED grow lights (PrimeGarden) with 16 hrs light for vegetative growth and 12 hrs light for flowering at 22°C.

### Pollen germination assay (PGA)

#### Pollen germination media (PGM)

The pollen germination assay was adapted from Schreiber & Dresselhau (2003) with some modification. The original protocol from Schreiber & Dresselhau (2003) employed 1% noble agar. In our study, we tested pollen germination media as liquid or combined with 1% agar. We found that liquid media resulted in better image acquisition and quantification of germination than solid media. During optimization of the Pollen Germination Assay (PGA), pollen concentrations of 0.1, 1 and 10 mg/mL were employed with the pollen diluted in PGM and incubated for 16 hr.

A 2X PGM contained the following: 10% sucrose (BIOSHOP), 0.005% H_3_BO_3_ (Sigma), 10 mM CaCl_2_ (BIOSHOP), 0.05 mM KH_2_PO_4_ (Merk) and 6% PEG 4000 (Fluka). After components were added to distilled H_2_O, heated on a stir plate for 10 mins at 70°C then filter sterilized. A 1X working solution was prepared fresh each day by diluting in distilled H_2_O.

#### Pollen collection and optimization of the PGA

Pollen was obtained from flowering male *Cannabis sativa* plants using a vacuum manifold method (Johnson-Brousseau and McCormick, 2004). For the standardized PGA, 10 mg of cannabis pollen was combined with 1 mL of freshly prepared 1X PGM and diluted to 0.1 mg/mL. 200 µL was then pipetted into a 24-well tissue culture plate (Flat Bottom Cell+, Sarstedt) and sealed with parafilm. Plates were incubated in the dark at 22°C for 16 hours and examined using an inverted light microscope (Zeiss Axio Observer Z1, Germany).

#### Imaging and germination assessment

Images were taken using phase contrast at 100x magnification. For each technical replicate, 8 images were taken to get an accurate representation of germination. Pollen germination percentages were calculated by dividing the number of germinating pollen grains by the total number of pollen grains. Germination percentages for each replicate represent the averages of the eight images.

### DAPI staining of cannabis pollen to decipher its bicellular or tricellular nature

Collected pollen was stained with DAPI (4′,6-diamidino-2-phenylindole) and imaged using an inverted fluorescent microscope (Zeiss Axio Observer Z1). The DAPI staining protocol was adapted from Backues et al. (Backues et al., 2010). Germinated pollen was suspended in pollen isolation buffer (PIB) containing 100 mM NaPO_4_(pH 7.5), 1 mM EDTA, 0.1% (v/v) Triton X-100 and 1 µg/ mL DAPI. A drop of solution was placed on a coverslip, incubated at room temperature for 5 minutes and viewed with the DAPI filter set. For DAPI staining of germinated pollen, pollen germination was performed as previously described, and staining was conducted with 1 µg/ mL DAPI after 16 hrs in PGM.

### Pollen collection from different development stages to assess the loss of pollen viability over time

Pollen was collected from male *Cannabis sativa* plants at different stages of the flowering development. The four stages of flowering were chosen according to the BBCH (Biologische Bundesantalt, Bundessortenamt and Chemische) scale adapted for cannabis (Mishchenko et al., 2017) and are listed as follows with the BBCH notation in brackets: Early (62), Mid (64), Mid-Late (65) and Late (67). Early stage (62) flowering was characterized by green mostly (80%) unopened anthers. By Mid stage (64) approximately 40% of anthers were open while Mid-Late was characterized by 50% opened. By Late stage (68) flowering was over and many of the flowers had turned yellow and fallen.

Following collection, an aliquot from each developmental stage was taken and used in the PGA for germination rate at time of collection (T0). The rest of the aliquots were stored in a 1.5 mL centrifuge tube at 4°C in the dark. After one week an aliquot from the different developmental stages was used for a PGA (T1), again after 2 weeks (T2) and again after 3 weeks (T3).

### Cryopreservation of pollen

Cannabis pollen submerged in Liquid Nitrogen (LN) without the use of any cryoprotectant or treatment will fail to germinate after the formation of ice crystals (Bajaj, 1987; Towill et al., 2000). Cannabis pollen was combined with cryoprotectants DMSO or glycerol diluted to concentrations of 10, 20, 30 and 60% and submerged in LN. Following 24 hrs in LN, pollen was removed and used in a PGA as previously described.

### Desiccation of pollen prior to cryopreservation

Cannabis pollen was combined with all purpose baked wheat flour (1:10 w/w) in a 1.5 mL centrifuge tube and desiccated at 5, 15 and 25 kPa for 20 or 40 mins. Following desiccation, the tube was placed in LN for four months, removed and placed at 22°C for 10 mins. The pollen/ wheat flour mix was then used for a PGA as previously described.

## Results and discussion

### Optimization of pollen germination for PGA

To obtain a representation of germination profile, a time-lapse of a pollen germination assay (PGA) was performed. We have observed the germination profile for 6 hrs with 30 min interval. The final germination was calculated after 16 hrs incubation. Germination started within 30 minutes with extending pollen tubes clearly visible (Figure 1 and Graphics Interchange Format (GIF) in supplementary information).

**Figure 1:**
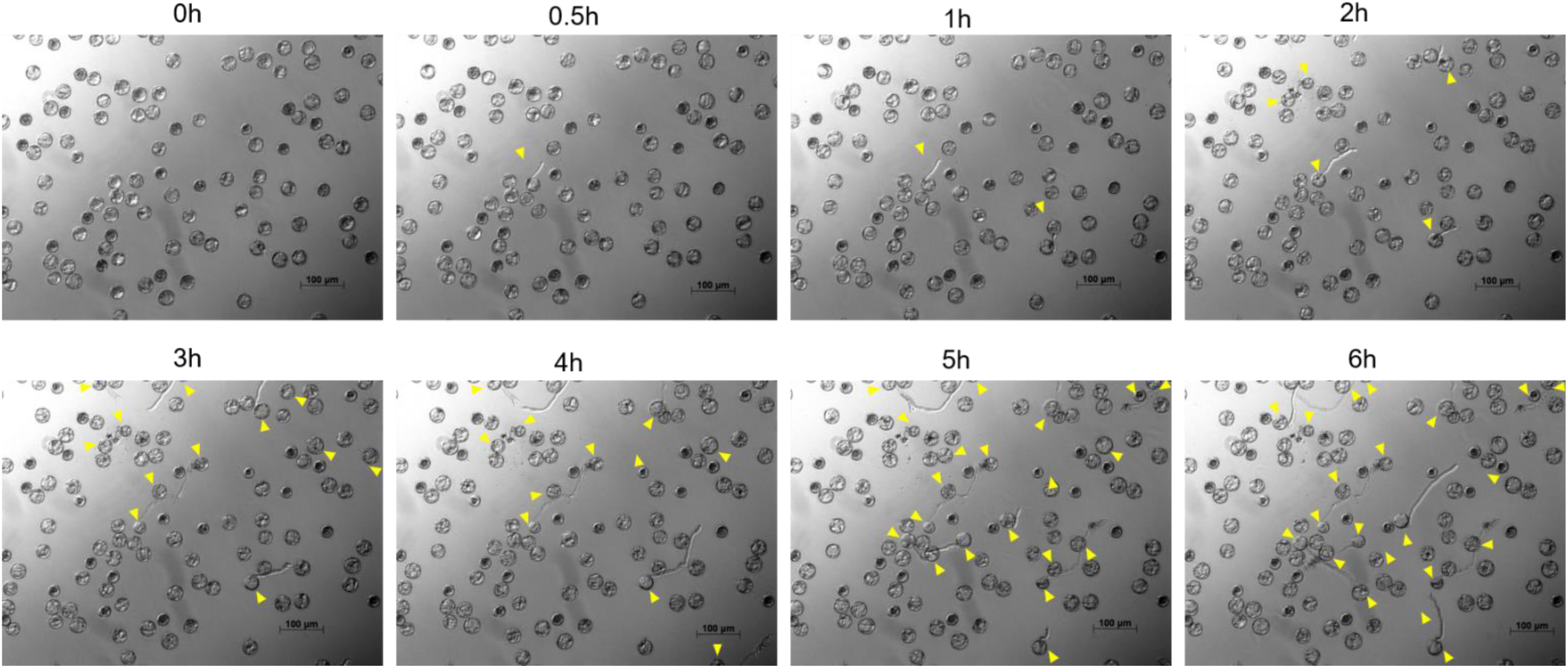
Representative photographs of cannabis pollen germination profile. Images taken at every 30-minute intervals for 6 hrs with germinating pollen grains indicated by the yellow arrows. Germination was started within 30 minutes with extending pollen tubes clearly visible. Images were acquired using an inverted fluorescent microscope (Zeiss Axio Observer Z1, Germany).

Collected cannabis pollen readily germinated in the Pollen Germination Media (PGM). PGM was tested both as a liquid and in combination with 1% agar. Germination rates were comparable in both tested media, however, pollen tubes were not as easily imaged under the microscope when germinated on agar (data not shown). For this reason, we opted for performing the PGA using liquid media. Of the different concentrations of pollen tested, 0.1 mg/mL provided the clearest imaging of germination, as higher concentrations resulted in crowding in the test well and reduced visibility (Figure 2). Additionally, in the highest density treatment, germination was adversely affected and made it difficult to accurately quantify germination percentage (Figure 2).

**Figure 2:**
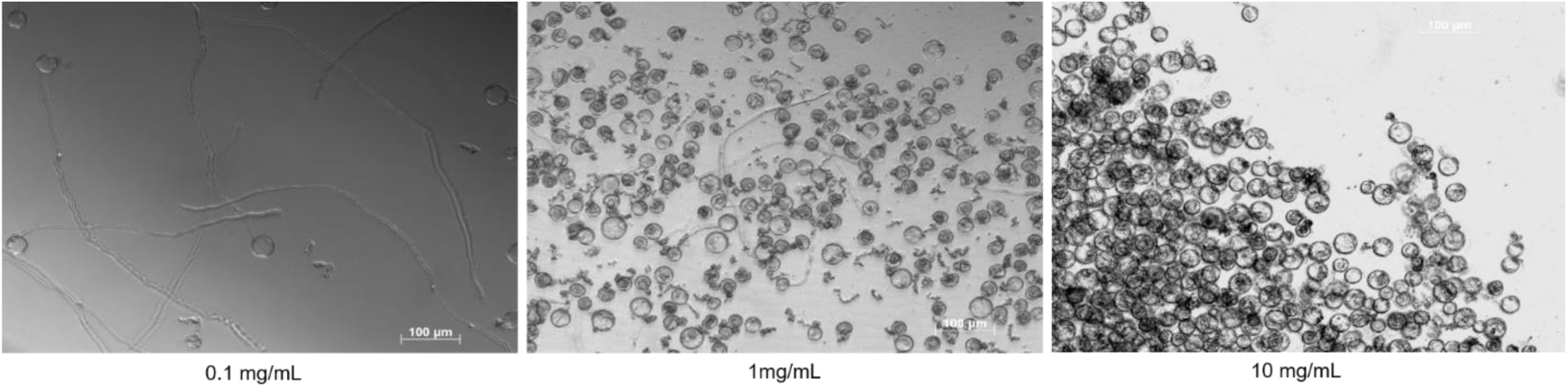
Optimization of Pollen Germination Assay. Cannabis pollen germination in PGM at concentrations of 0.1, 1 and 10 mg/ mL. Images were acquired after 16 hours using an inverted fluorescent microscope (Zeiss Axio Observer Z1, Germany).

### Pollen collected from different principal growth stages exhibits different longevity

To establish how cannabis pollen germination rates change over time, we tested the pollen in a pollen germination assay after storage at 4°C. Because pollen collected from different principal growth stages may affect germination rates, we collected pollen from males at different points during floral development to cover the entirety of anthesis.

We compared the loss of viability of cannabis pollen collected from the four different points during flower development over the course of 21 days. The rate of germination at T0 was comparable amongst the different developmental stages with maximum 50% germination in Mid-Late (65) stage, with all stages losing viability after only one week at 4°C storage (Figure 3). While loss of pollen viability was apparent from pollen collected from all time points, pollen collected from Mid flowering stage (64) appeared to retain viability the longest with 22% of pollen grains successfully germinating after 21 days of 4°C storage (Figure 3). This may indicate that an optimal growth stage for pollen collection is around developmental stage (64), whereas the loss of pollen viability may begin while the pollen is still present in the anthers. Pollen collected earlier, at developmental stage 62, may have not fully matured resulting in a lower germination percentage (Figure 3).

**Figure 3:**
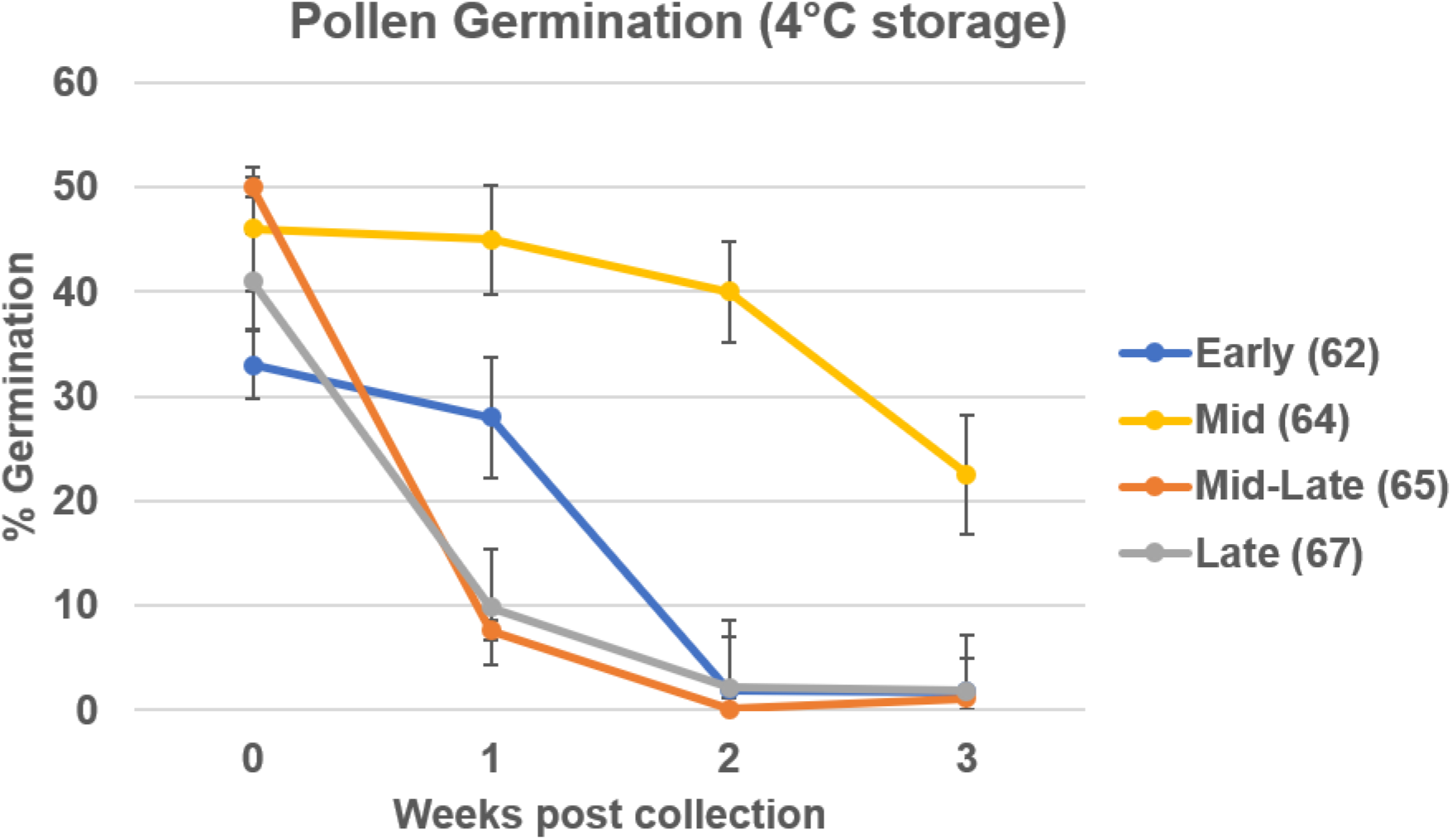
Loss of pollen viability over time. Pollens was harvested from plants at four different developmental time points then stored at 4°C for one to three weeks. Viability was determined via pollen germination assay. Data are shown as average ±SD.

### DAPI staining revealed bicellular nature of cannabis pollen

While the fluorescein diacetate (FDA) stain is routinely used for viability tests, it is not ideal for visualizing the nuclei in pollen cells. In order to establish whether cannabis was bicellular or tricellular, we performed a DAPI stain on germinating cannabis pollen. Prior to pollen tube germination, the brighter, more compact sperm nucleus and the diffuse vegetative nucleus were visible (Figure 4 A-B). The brighter staining in the sperm nucleus represents the more condensed state of chromatin compared to the more transcriptionally active vegetative nucleus. Following pollen tube germination, both sperm nuclei are clearly visible as they descend the pollen tube (Figure 4 C). This suggests that cannabis releases sexually immature pollen grains, with the second mitosis event occurring after pollen tube germination.

**Figure 4:**
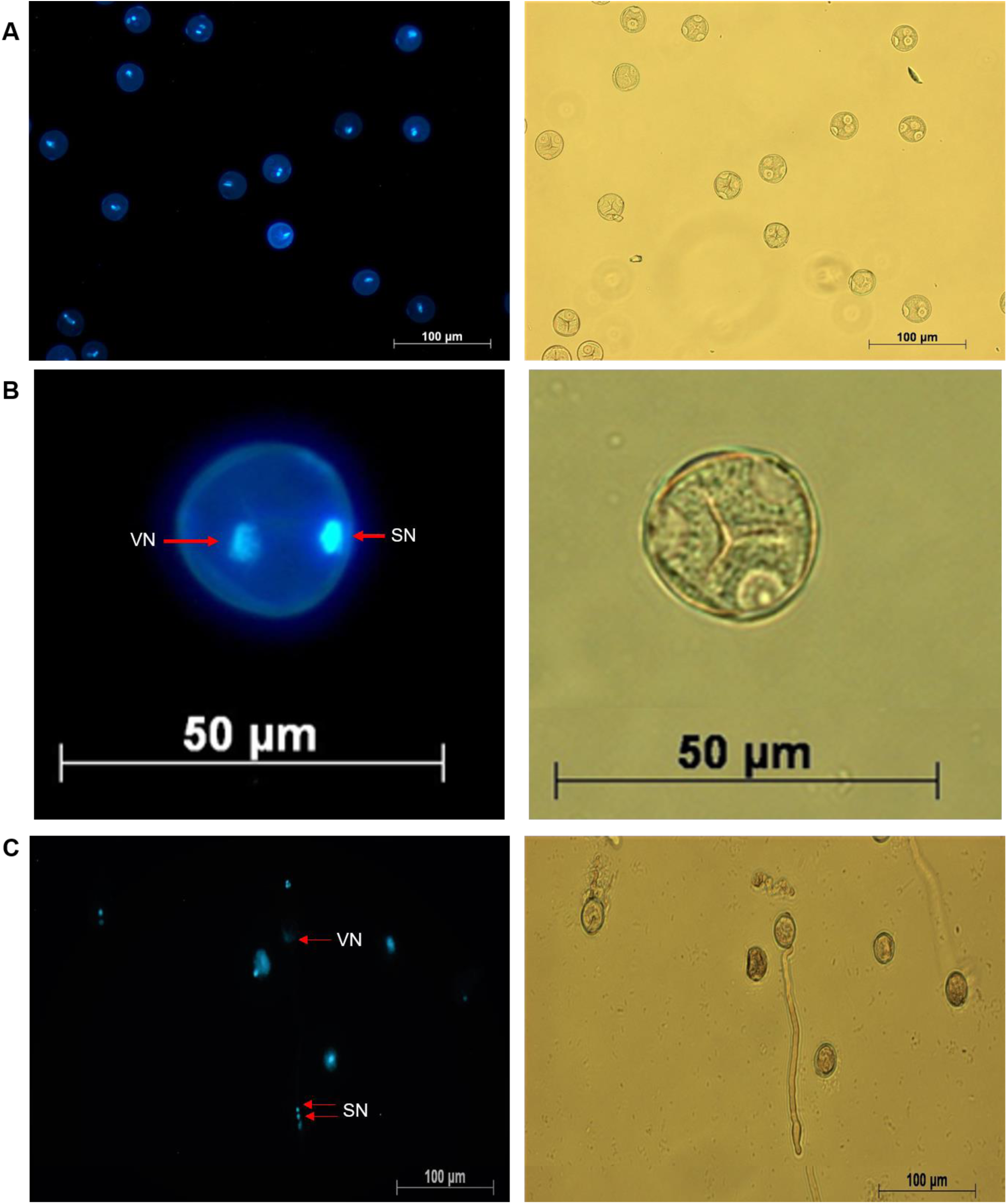
Visualization of cannabis pollen at different stages of germination using DAPI staining. Both the sperm (SN) and the vegetative nuclei (VN) are visible at the bicellular stage prior to pollen tube germination (A,B). Image C represents a germinated cannabis pollen cell with the migrating two sperm nuclei indicated by the red arrows. Images on right side panel represent the phase contrast view of respective left image. Images were acquired using an inverted fluorescent microscope (Zeiss Axio Observer Z1, Germany).

### Development of a cryopreservation method for cannabis pollen

Pollen cryopreservation has been attempted in a variety of agriculturally and medicinally important plant species for the preservation of elite germplasm. Numerous studies have reported the data on pollen viability under various storage conditions (Towill, 1985; Engelmann and Takagi, 2000). While the interaction between pollen water content and viability is complex, it is understood that an optimum water content is necessary for longevity (Engelmann and Takagi, 2000). Generally, longevity is increased by lowering the temperature and moisture content. Some reports indicate a moisture optima of 15%, while higher water concentrations (above 30%) may result in rapid deterioration (Buitink et al., 1998). Liquid Nitrogen (LN; −196 °C) is routinely used for cryogenic storage, as it is relatively cheap, safe and at a temperature where enzymatic and chemical reactions do not cause biological deterioration (Kartha, 1985). Cannabis pollen stored in LN without prior desiccation failed to germinate (data not shown). Pollen cells with high moisture levels do not survive cryogenic storage, presumably due to intracellular ice formation (Engelmann and Takagi, 2000). Therefore, pollen cells need to be dried within a range where no freezable water exists without succumbing to desiccation injury. For pollen desiccation, we tested a vacuum desiccation at various pressures (5, 15 or 25 kPa for either 20 or 40 minutes). When pollen was desiccated prior to storage in LN, it failed to germinate (data not shown). We therefore assumed that desiccation alone may not be sufficient for preservation of pollen viability via cryopreservation. In addition to desiccation, we also tested other cellular cryoprotectants such as DMSO and glycerol which may improve cell survival after cryogenic storage (Pritchard, 1989). Desiccated cannabis pollen combined with a 10, 20, 30 or 60% DMSO or glycerol solution prior to being stored in LN for 24 hours had a 0% germination rate (data not shown).

Baked wheat flower has been previously suggested as a possible cryoprotectant for long term pollen storage (Yi et al., 2003). To test this hypothesis, cannabis pollen was desiccated and combined with baked wheat flour. Vacuum desiccation at a lower pressure of 5 kPa for the longest interval for 40 minutes, resulted in the highest germination rate after storage in LN after 24 hours (Figure 5). Higher pressures did not result in pollen germination, as the cells may have been compromised during the drying process (data not shown). This treatment was used for subsequent preservation experiments where we achieved an average germination rate of 16.4% compared to the non-LN control (control was subjected to same desiccation combined with whole baked wheat flour) of 26.3 % (Figure 6). Desiccation itself cause around 50% reduction in germination as compared to untreated freshly harvested pollen (Figure 3 & 6). There was a 10% reduction observed in germination after 24 hours in LN as compared to non-LN control (control was subjected to same desiccation combined with whole baked wheat flour). Desiccated cannabis pollen was kept in LN for four months to test long term storage and was found to have an average germination rate of 14.6 % (Figure 6), suggesting long term storage is a possibility under appropriate conditions. To confirm *in planta* viability of the treated cannabis pollen, the pollen/ wheat flour mix was removed from LN and applied to flowering female cannabis plants. The pollination resulted in successful seed formation in all the flowers receiving treated pollen.

**Figure 5:**
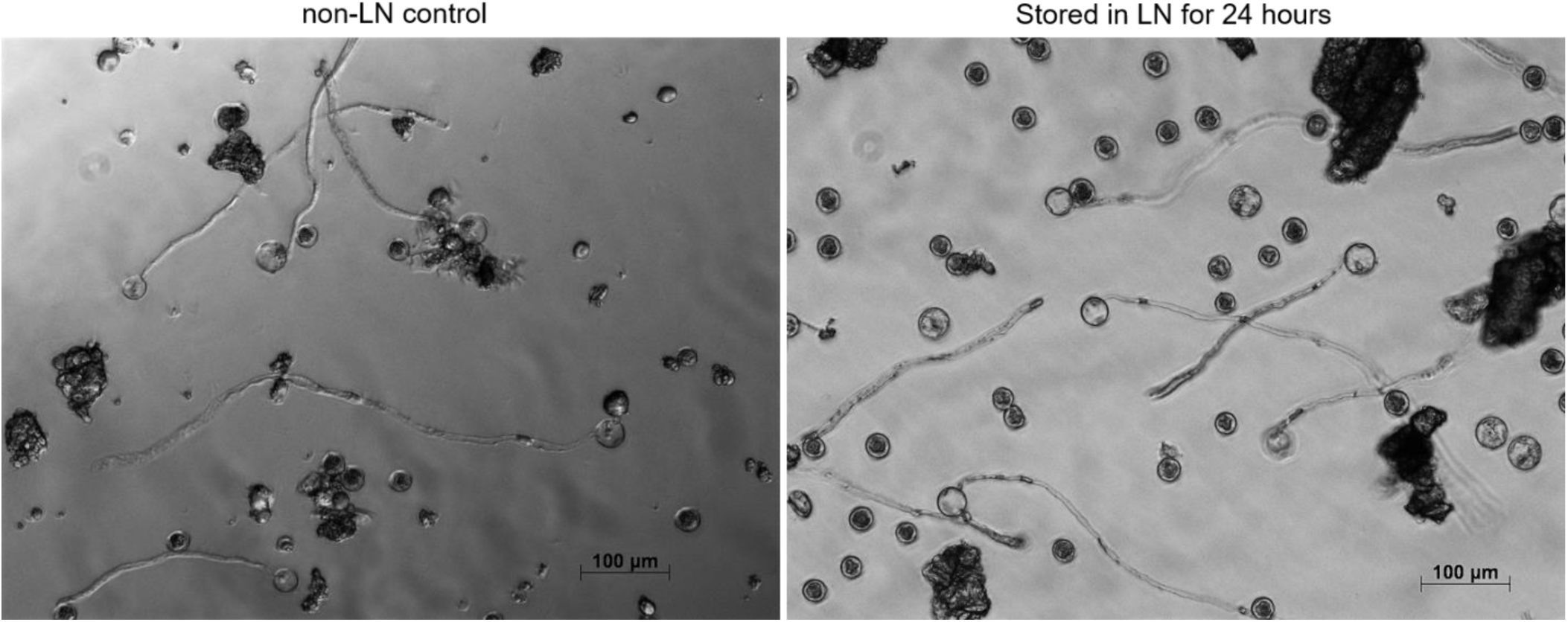
Representative photographs from pollen germination assay (PGA) of pollen stored for 24 hrs in liquid nitrogen. Desiccated cannabis pollen mixed with 1:10 wheat flour and stored in liquid Nitrogen. Non-LN control (control was subjected to same desiccation combined with whole baked wheat flour). Pollen flour mix was diluted to 0.1 mg/mL and used for PGA.

**Figure 6:**
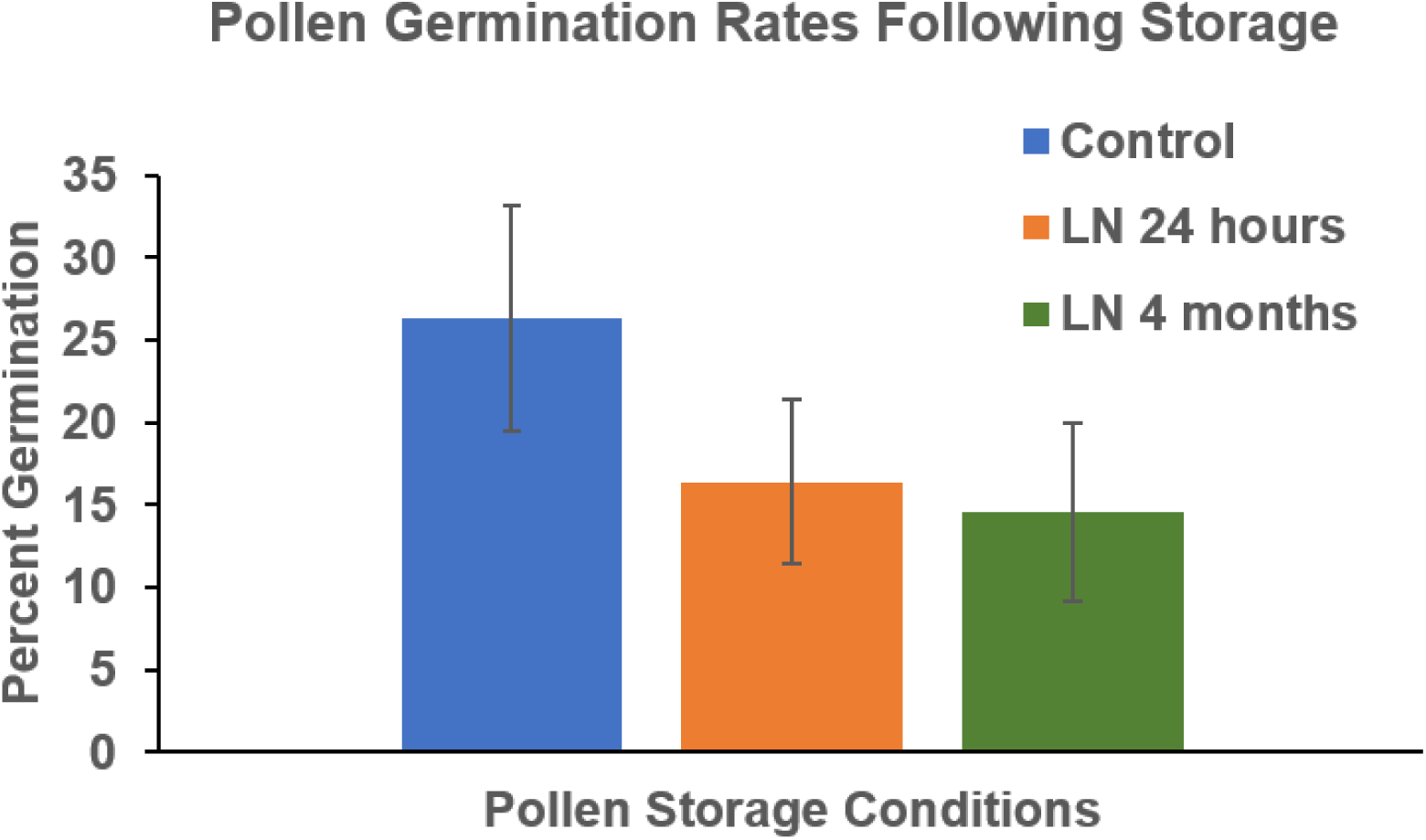
Comparison of pollen germination efficiency between pollen stored in liquid nitrogen for 24 hours, 4 months and non-LN control (control was subjected to same desiccation combined with whole baked wheat flour). Data are shown as average ±SD.

There are several reports on pollen long-term cryopreservation including one-year viability (*Allium* sp., (Kanazawa et al., 1992); *Juglans nigra*, (Luza and Polito, 1988); *Diospyros khaki* (WAKISAKA, 1964)), two years viability (*Jojoba*, (Lee et al., 1985); *hop*, (Haunold and Stanwood, 1985)), and five or more years survival (*Vitis vinifera L*., (Ganeshan and Alexander, 1990); tomato and eggplant, (Alexander and Ganeshan, 1989); Maize, (Barnabas, 1994); Gladiolus (Rajasekharan et al., 1994)). Our cryopreservation method resulted in a 10% germination loss after 24 hrs of LN storage, whereas a 4-month storage in LN further reduced the germination rate only by an additional 1.8% (Figure 6). Similar to our report, Hamzah and Chan (1986), also suggested viability declines over a relatively short time (Hamzah and Chan, 1996). *Hevea* pollen exhibited a decline from 20% in vitro germination after 1 month to 2% after 5 months of storage in LN. Some pine and spruce pollen stored in LN also showed a decline in viability over a 24-month period (Lanteri et al., 1993). Cryopreservation of maize, lily (Nath and Anderson, 1975) and wheat pollen (Andreica et al., 1988) also exhibited a decline in viability during cryopreservation. Overall, our results suggest that periodic viability testing of cryopreserved pollen is required to assure the future use of stored pollen in breeding.

In conclusion, we have standardized a simple assay for quickly assessing pollen germination in *Cannabis sativa*. Through the use of DAPI staining on germinating pollen cells, we were able to track the migration of sperm nuclei descending the pollen tube. This indicates that *Cannabis sativa* releases pollen in a bicellular state, where the second mitosis event occurs after pollen tube germination. By using our PGA, we have demonstrated the loss of pollen viability over time when stored at 4°C, and suggested an optimal time during flower development for pollen collection to maximize longevity during storage. Finally, we have provided an easy protocol for cryopreservation using desiccation combined with baked wheat flour and subsequent long-term storage of cannabis pollen in liquid nitrogen.

## Acknowledgements

We thank Natural Sciences and Engineering Research Council of Canada (NSERC) and MITACS for funding our work.

## Competing interests

The authors declare that they have no competing interests.

